# RNAdvisor: a comprehensive benchmarking tool for the measure and prediction of RNA structural model quality

**DOI:** 10.1101/2023.11.14.567018

**Authors:** Clément Bernard, Guillaume Postic, Sahar Ghannay, Fariza Tahi

## Abstract

RNA is a complex macromolecule that plays central roles in the cell. While it is well-known that its structure is directly related to its functions, understanding and predicting RNA structures is challenging. Assessing the real or predictive quality of a structure is also at stake with the complex 3D possible conformations of RNAs. Metrics have been developed to measure model quality while scoring functions aim at assigning quality to guide the discrimination of structures without a known and solved reference. Throughout the years, many metrics and scoring functions have been developed, and no unique assessment is used nowadays. Each developed assessment method has its specificity and might be complementary to understanding structure quality. Therefore, to evaluate RNA 3D structure predictions, it would be important to calculate different metrics and/or scoring functions. For this purpose, we developed RNAdvisor, a comprehensive automated software that integrates and enhances the accessibility of existing metrics and scoring functions. In this paper, we present our RNAdvisor tool, as well as state-of-the-art existing metrics, scoring functions and a set of benchmarks we conducted for evaluating them. Source code is freely available on the EvryRNA platform: https://evryrna.ibisc.univ-evry.fr.

## Introduction

The various types of non-coding RNA molecules exert their biological functions either by base-pairing mechanisms or through their three-dimensional structure. As for proteins, experiments for determining the spatial conformation of RNA chains are costly, which has led to the development of computational methods for predicting the biologically active (“native”) fold from the sole ribonucleic acid sequence (1–5). However, these methods dedicated to RNA have yet to reach the same accuracy as their protein-specific counterparts, such as AlphaFold2 (6) or ESMFold (7). The fact that the native structures available in the Protein Data Bank (PDB) are far less numerous and diverse for RNA than for protein molecules mainly explains this slower rate of progress in solving the RNA folding problem. Nevertheless, current efforts are put forth to overcome this lack of training data.

The quality of a structural model is defined by its “nativity” or native-like character, i.e. how close it is to the na-tive fold of the same RNA sequence. Therefore, evaluating the performance of a predictive method requires measuring the similarity between the 3D models it generates and the corresponding native RNA structure. For this model quality measure, multiple metrics have been proposed throughout the years. Some were directly transposed from the study of protein structures, such as the root-mean-square deviation of atomic positions (RMSD) or the template modeling score (TM-score) (8). Others, such as interaction network fidelity (INF) (9) or mean of circular quantities (MCQ) (10), have been created to take into account the specificities of RNA 3D structures, in particular their greater flexibility.

For predicting RNA fold from the sequence, algorithms explore the conformational space through different strategies (11–15). This produces a certain number of predicted RNA structures that must be ranked. In a real-case scenario where the native structure is not known, such a ranking requires computing relative quality predictions for all the generated models. For this purpose, different scoring functions, also called model quality assessment programs (MQAP), have been released (16–19). Evaluation of these scoring functions is usually done with near-native structures called decoys, which are disturbed native structures that play the roles of predicted RNA structures for predictive models.

An ideal score for predicting model quality would correlate with the Gibbs free energy change (Δ*G*) of the RNA folding process, as the native structure is the one with the most negative Δ*G*_folding_. However, calorimetric data are not available for the unfolded states, so the thermodynamic relevance of the MQAP scores cannot be evaluated directly. The predicted quality scores are diverse and computed through different approaches, which raises the question of the equivalence between these metrics for representing the ground truth, i.e. the nativity of the model. In case where they actually represent different aspects of the nativity, a subsequent question regards the dependence of the MQAP’s accuracy on these different model-to-native similarity measures. To facilitate the calculation of different metrics and scoring functions for a better evaluation of RNA 3D structure predictions, we developed a computational tool called RNAdvisor. RNAdvisor is an open-source tool that integrates all available codes of state-of-the-art metrics and scoring functions.

In this paper, we bring a comprehensive interpretation of quality measurement and model quality assessment of RNA 3D structures. We describe our RNAdvisor tool, before presenting the benchmarks we conducted thanks to RNAdvisor. Different benchmarks were carried out, considering three datasets available online. We evaluated the scoring functions, and measured the relationship between the different metrics, and between the scoring functions and metrics. We also measured the running time, as well as the CO_2_ equivalent consumption. All these benchmarks are reproducible, open-source, and accessible at https://github.com/EvryRNA/rnadvisor_results.

### State-of-the-art metrics

To assess whether a predicted RNA tertiary structure is close to its native fold, multiple metrics have been developed. Quality measurement can be general, telling how well the prediction falls into the global conformation. Other metrics, inspired by protein metrics, consider the alignment of structures to evaluate a predicted structure. Nevertheless, proteins and RNAs have differences that limit the adaptation of protein metrics to RNAs. One of the significant differences lies in folding stabilization, where RNA structure is maintained by base pairing and base stacking, while hydrogen interactions in the skeleton support protein structure. Therefore, metrics have been developed to fit the RNA specificities, considering the different types of interactions.

A summary of the state-of-the-art metrics is provided in Table 1.

**Table 1.** Summary of the state-of-the-art metrics used for assessing the quality of
predicted RNA 3D structures compared to a reference.

| Metric | Granularity |
| --- | --- |
| **General metrics** |  |
| RMSD | Atom deviation |
| CLASH (20) | Steric clashes |
| εRMSD (19) | Relative distance and orientation |
| **Protein-inspired** |  |
| TM-score (8) | Normalised atom deviation |
| GDT-TS (21) | Count of superimposed residues |
| CAD-score (22) | Contact-area |
| IDDT (23) | Interatomic distance differences |
| **RNA-oriented** |  |
| P-VALUE (24) | Non-randomness assessment |
| INF, DI (9) | Key interactions accuracy |
| MCQ (10) | Angles dissimilarity |

#### A. General metrics

The general metrics give an overall idea of the quality of a prediction. They are usually based on an overall distance averaged throughout the structure. The most used metric is the RMSD, which gives an overall predicted model evaluation. An improvement of this metric was proposed with *ϵ*RMSD (19), which incorporates RNA features. On the other hand, the CLASH score (20) assesses the overlaps of atoms and doesn’t consider the atom deviations compared to the previously mentioned metrics. The main advantage of general metrics is the quick overview of the nativity of the structure. It gives a unique value, an averaged similarity score over the reference. The RMSD is almost always used as a criterion to assess the quality of a computational approach in a database. Nonetheless, it is limited in explaining the limits of a prediction. A high dissimilarity in a small region would highly bias the RMSD value. The CLASH score is more used as an assessment of possible conformation. An almost native structure would have a very low CLASH score, while a low CLASH score structure doesn’t necessarily mean a native structure. Finally, the *ϵ*RMSD tries to add relative base arrangement to the atomic distance deviation to incorporate RNA structural features.

#### B. Protein-inspired metrics

Although proteins and RNAs are different molecules, conformational folding shares few characteristics. A higher proportion of solved protein structures makes developing ap-proaches easier. Consequently, protein metrics have been studied and widely used, especially in the CASP competition. One of the known metrics is the TM-score (8), which adds distance normalization to a classic RMSD. Given aligned structures, the GDT-TS (21) computes superimpositions with different distance cutoffs. Another approach, with CAD-score (22), is using a contact-area function to assess differences. The lDDT (23) score was created to quantify the model quality on the level of the residue’s environment, where local atomic interactions are considered to obtain a robust metric. The conception of those metrics is not restricted to proteins and can be adapted to RNA sequences. The proteins-based metrics adapted to RNA molecules can give a general overview of predicted structures. While the TM-score avoids the increase of deviation score if the sequence increases, it is still limited to a general assessment. CAD-score and GDT-TS try to incorporate local superimposition, but it would still suffer from the lack of local information.

#### C. RNA-oriented metrics

RNAs are unique molecules with a tertiary conformation maintained by base pairing and base stacking. The torsion angles that describe each nucleotide, such as the approximated pseudo-torsion, can be used to best assess the nativity of a structure compared to a solved structure. RNA also has well-defined pairing patterns, where a base interacts with each other. These interactions are very specific, and general metrics or scores inspired by proteins can not integrate them. That is why multiple metrics like INF (9) (for base pairing patterns) or MCQ (10) (for torsion angles) have been developed to allow the integration of RNA structural specificities. The INF score can be specific to base-pairing interactions (INF_*bp*_), the base-stacking interactions (INF_*stack*_), or consider both (INF_*all*_). To include both RMSD and INF advantages, the deformation index (DI) (9) has been developed as the quotient of RMSD by INF. Another metric is the P-VALUE (24), which assesses the validity of a prediction: it describes if a prediction is better than a random prediction. Metrics specific to RNA have the advantage of considering specificities that are major parts of RNA 3D structure stabilization. The metrics have a more concrete meaning and could help the comprehension of a failing prediction. For instance, a bad *INF*_*bp*_ (value near 0) value would mean a failing in base-pairing interactions, whereas a bad RMSD (high value) does not provide this information (and could also be biased by a local misprediction). RNA-oriented metrics remain complementary: INF and MCQ scores describe different structure characteristics. Those RNA-oriented metrics can be added to general and protein-based evaluation metrics for a near-complete assessment of predicted structures.

No unique metric can assess structure quality. Each metric has a different particularity that can complement other metrics. While the RMSD and INF are widely used in the community, their efficiency remains limited with real-world RNAs.

A complete description of the different metrics is provided in Supplementary Materials. It also details the different implementation languages, as well as the source codes.

### State-of-the-art scoring functions

The nativity of RNA molecules can be computed by dissimilarity metrics but requires having a known solved reference structure. This requirement is challenging as the number of solved structures is low. Furthermore, computational methods usually predict multiple conformations that need to be ranked. The relative quality prediction can not rely on a known solve structure. The adaptation of the free energy of the structure has become a standard in the ranking, filtering and confidence assessment of structures. These predictive quality measurements are knowledge-based approaches that rely on statistical potentials. With the recent success of AlphaFold2 (6), new approaches employ deep learning methods for quality predictions of RNA structures.

A summary of the different state-of-the-art scoring functions is provided in Table 2.

**Table 2.** Summary of the scoring functions used for assessing the nativity confidence of RNA 3D structures.

| Score | Granularity |
| --- | --- |
| **Knowledge-based** |  |
| RASP (16) | Pairwise-distances |
| eSCORE (19) | Relative positions |
| 3dRNAScore (33) | Distance and dihedral angles |
| DFIRE-RNA (18) | Pairwise-distances |
| rsRNASP (17) | Short and long pairwise distances |
| **Deep learning** |  |
| RNA3DCNN (34) | Atom grid |
| ARES (35) | Atom position and chemical type |

#### D. Knowledge-based scoring functions

Prediction-based methods like NAST (11), HiRE-RNA (4) or SimRNA (12) use an adaptation of the free energy in their discriminative phase. A common approach uses knowledge-based statistical potentials considering structures to create a quality measurement score. It has been proven to work well for proteins, such as the one used in AlphaFold 2 (6). These potentials are said to be derived from Boltzmann formulations. They rely on a comparison with non-native base pair interactions, known as a reference state. The reference state should ideally come from a set of non-redundant decoy conformations where no interactions between atoms appear. Unfortunately, no ideal dataset exists (25), but approximations of reference states have been proposed through the years (26– 31). Adaptation to RNA has been studied (32) and remains limited by the lack of a large and representative RNA dataset.

Most knowledge-based approaches to assess RNA structure nativity employ an all-atom distance potential and use averaging reference states, like 3dRNAScore (33) or RASP (16). The challenge is to find good structural features that consider RNA conformational specificities to distinguish native and non-native folding. Methods like *E*SCORE (19) or DFIRE-RNA (18) consider relative orientation to incorporate RNA flexibility. Short and long-range interactions are considered differently with different reference states in the new potential rsRNASP (17). The main limitation of knowledge-based scoring functions is the lack of a dataset of reference state decoys.

#### E. Deep learning scoring functions

With the recent success of AlphaFold2 (6) and its deep architecture, MQAP scores have been developed like RNA3DCNN (34) or ARES (35). They input different characteristics like chemical type or atom position. They use available native conformations to learn a score without explicitly using a reference state. The objective is an RMSD-like metric, meaning that the network learns atom deviation properties to assess structure predictive quality. The architecture is based on a neural network with either convolutional layers or graph neural networks. They rely on decoy datasets generated by either FARFAR 2 (2) for ARES, or relaxed structures by molecular dynamics from PDB for RNA3DCNN. ARES and RNA3DCNN scoring functions remain limited by the current deep learning drawbacks: the lack of interpretability and the need for large datasets. As the number of solved RNA 3D structures is low, deep learning approaches could easily lack generalization to new unseen structures. Datasets considered are biased by either the chosen model creating decoys or the method to relax structures.

No ideal scoring function exists, and the available scores can also be complementary: one score can weigh more dihedral angles, whereas the other could consider chemical types. As no ideal metric exists, some scoring functions could be more linked to a given metric, making the ranking more difficult.

A complete description of the different scoring functions is available in Supplementary Materials, as well as the different implementation languages and the source codes.

### RNAdvisor tool

As the number of available RNA 3D structures increases, assessing the nativity of predicted structures becomes crucial. Numerous families still have unsolved structures in the PDB, but they might be available in the following years. Assessing and understanding the limits of predicted methods for the available and nearly available RNA 3D structures is essential. As discussed previously, no perfect metrics can discriminate between native-like and wrong-predicted structures. The same goes for the scoring functions. Each metric or scoring function has its specificity and could complement the understanding of RNA conformation. Nonetheless, metrics and scoring functions have been developed for years by different researchers in different programming languages, making their use difficult. Some web servers, like RNA-tools (36), can compute RMSD or INF scores. It was introduced after RNAPuzzles (37), a collective challenge to evaluate predicted 3D RNA structures. Web servers might be helpful for discrete tests but can not be used to automate the evaluation process. As the number of RNAs is growing, we can not rely on web servers to check each of the predicted structures. Automating the computation of scoring functions is even more crucial as they are widely used for sampling procedures.

We developed a tool called RNAdvisor, that enables the computation of all the available state-of-the-art metrics and scoring functions in one command line. It integrates eleven metrics and four existing scoring functions from nine standalone codes, as shown in Figure 1. We omitted 3dRNAscore because we could not get the source code. We failed to run the ARES code, and RNA3DCNN had bad results compared to the published ones, so we decided not to include it.

**Figure 1.**
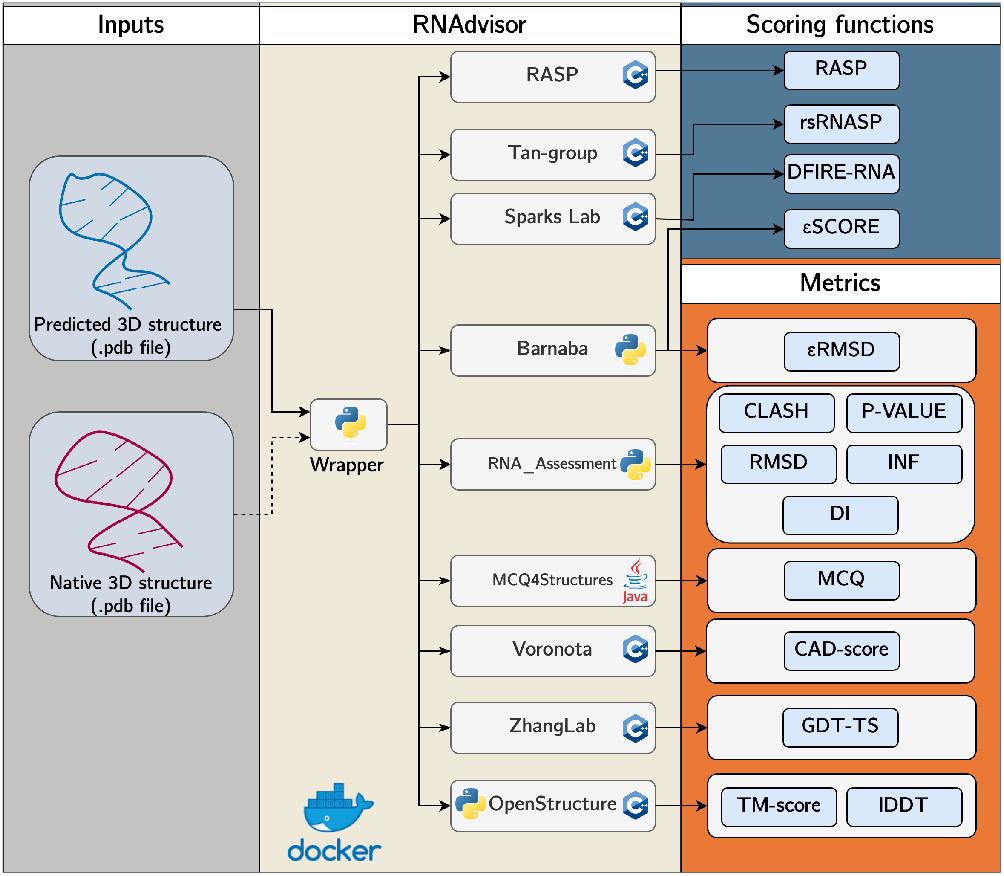
Schema of RNAdvisor tool: a wrapper code that gathers open-access libraries for assessment of RNA 3D structures in one interface. It is wrapped in a Docker image to emancipate laborious installation processes. A user can input an RNA 3D structure with the reference structure, and it will compute each metric and scoring function using the different integrated software.

Our tool uses coding best practices like DevOps library, named Docker (38) to emancipate the dependency of OS. All the installation needed by each library is already done and easily accessible.

### Benchmark

To evaluate the performance of a scoring function, a common practice (16, 17, 34, 35) is to compare the rank obtained by the native structure in a set of decoys. In this section, we first describe the three datasets of decoys used for the experimentation, followed by a study of the link between existing metrics. Then, we examine the performance of scoring functions, followed by a study of their correlation with metrics. We finally provide a benchmark of computation time and CO_2_ emissions for scoring function and metrics.

#### F. Datasets

We used three datasets, named Test Set I, Test Set II and Test Set III, to assess the relations between scoring functions and metrics. The first two datasets have decoys generated by two distinct strategies widely used to compare scoring functions (16, 17, 34, 35). The last dataset is a real-case scenario where 3D structures from different model predictions should be ranked by nativity.

**Test Set I**^1^ is composed of 85 RNAs with decoys generated by MODELLER (39), a predictive model that is used to create decoys with different set of parameters. It uses Gaussian restraints for atom distances and dihedral angles, leading to 500 decoy structures for each RNA. The decoys are close to the native structures as only minor changes are made in the decoy creation.

**Test Set II**^2^ corresponds to the prediction-models (PM) subset from rsRNASP (17). It consists of 20 non-redundant single-stranded RNAs with decoy structures generated by four RNA 3D models (10 per model): FARFAR 2 (2), RNA-Composer (40), SimRNA (12) and 3dRNAv2.0 (41). It leads to 20 RNAs with 40 decoy structures for each native RNA. The created decoys are less close to the native structure as they use predicted models to create the decoys.

**Test Set III**^3^ is the RNA-Puzzles_standardized dataset. It comes from the competition that reproduces the protein CASP challenge for RNA: RNA-Puzzles (37). It contains 21 RNAs and dozens of decoy structures for each RNA. It is commonly used as the most realistic test set to assess the generalization properties of models. The decoys are not all close to the native structure.

#### G. Evaluation metrics

Identifying native structures from non-native or near-native is a property required by scoring functions. To assess the quality of a given scoring function, we used the Pearson correlation coefficient (PCC) and the enrichment score (ES). The PCC is computed between the ranked structures based on scoring functions and structures ranked by metrics and is defined as:

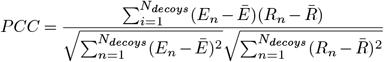

where *E*_*n*_ is the energy of the *n*th structure, and *R*_*n*_ the metric of the *n*th structure. PCC ranges from 0 to 1, where a PCC of 1 means the relationship between metric and energy is completely linear. The enrichment score (ES) is defined as:

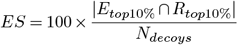

where |*Etop*10% ^∩^ *Rtop*10%| is the number of common structures from the top 10% of structures (measured by the metric) and the top 10% of structures with the lowest scoring function. ES ranges between 0 and 10 (perfect scoring). An enrichment score below 1 means a bad score and a value equal to 1 means a random prediction.

### H. Results

#### H.1. Metrics relationship

Each metric has its specificity and gives a result based on different assumptions: base interactions, angle conservation, atom distance deviation, etc. Metrics may be redundant with one other. We compared the PCC and ES between each computed metric averaged over the three datasets used. The results are shown in Figure 2. Details for each dataset are also provided in Supplementary file.

**Figure 2.**
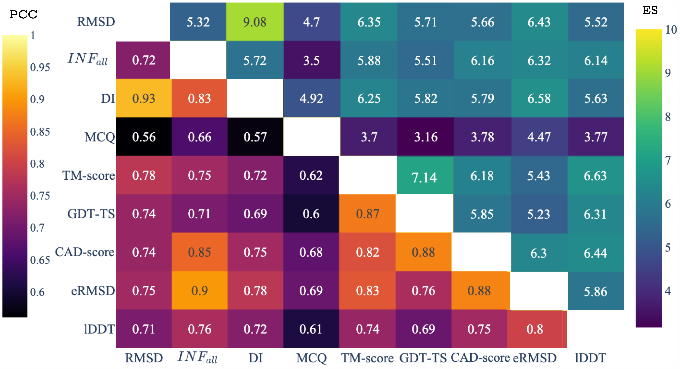
ES and PCC scores for each metric averaged over the three test datasets. The lower half of the matrix represents the PCC, while the upper half corresponds to the ES score. The diagonal has a PCC of 1 and ES of 10.

RMSD has a high correlation with DI, which is not relevant as DI is composed of both RMSD and INF metrics. RMSD correlates with *ϵ*RMSD in terms of ES and PCC (6.43 and 0.75, respectively) while being related TM-score (ES of 6.35 and PCC of 0.78). As *ϵ*RMSD tries to improve the classic RMSD and TM-score adds a normalisation, the correlation makes sense. INF metric highly correlates with ES and PCC with CAD-score and *ϵ*RMSD (ES of 6.16 and 6.32 and PCC of 0.85 and 0.9, respectively). DI is also linked to *ϵ*RMSD with an ES of 6.58 and PCC of 0.78. As the *ϵ*RMSD is an improved RMSD that includes RNA structure specificities, it makes sense that it is correlated to the DI metric as it includes both RMSD and RNA-specific INF metrics. MCQ is the only metric systematically less related to the other metrics. The angle consideration is not mainly included in other metrics computation, which could explain this behaviour. Nonetheless, MCQ has a high correlation for Test Set I with the other metrics (shown in the Supplementary file). It means that for near-native decoys, MCQ behaves like most other metrics, whereas with real-world prediction structures, it is uncorrelated to others. Near-native decoys might keep structural conformations and thus angle conservation, which is not true for the structures from Test Set I and Test Set II. TM-score is connected with another proteinbased GDT-TS metric with an ES of 7.14 and PCC of 0.87. Finally, lDDT metric is linked to TM-score (ES of 6.63 and PCC of 0.74), CAD-score (ES of 6.44 and PCC of 0.75) and INF_*all*_ (ES of 6.14 and PCC of 0.76). As the lDDT metric incorporates interatomic distance information, this is retrieved in the CAD-score and in the normalised atom deviation of the TM-score.

We can conclude that the MCQ metric is highly uncorre-lated to the others (for Test Set II and III), while the TM-score and GDT-TS seem very dependent. INF_*all*_ discriminate decoys with the same behavior as the CAD-score. *ϵ*RMSD is linked to INF and thus CAD-score (as they are correlated) and DI. Their correlations are not perfect (no ES of 10 or PCC of 1), meaning that every metric can help assess predicted model quality.

#### H.2. Scoring function ranking

The aptitude of a scoring function to classify native and near-native structures is essential for developing models. Table 3 shows the number of native structures with the lowest scoring function value among the decoys. RASP performs well for near-native structures (Test Set I) by finding 83 out of 85, but fails for the other datasets. *E*SCORE and DFIRE-RNA have almost identical results, with 99 and 103 found structures out of 126 overall. rsRNASP does not perform as well as the other scoring functions for Test Set I but outperforms them for the two other datasets. It leads the overall native structures found with 111 out of 126. Details of the average rank of the native structure for each dataset are provided in Supplementary file.

**Table 3.**
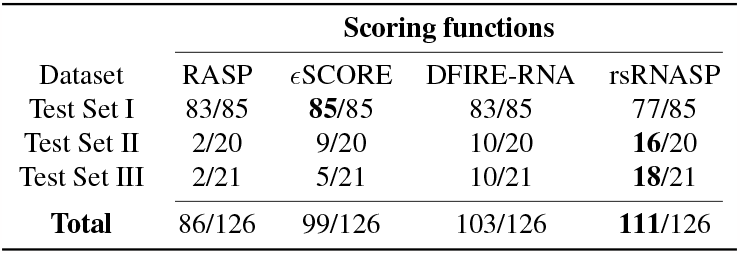
Number of native structures found with the lowest score for each dataset. It corresponds to the number of times the native structure has the lowest scoring function value among the decoys.

rsRNASP seems the best scoring function for ranking and finding the native structure, followed by DFIRE-RNA and *E*SCORE. rsRNASP is less accurate for close decoys (represented by Test Set I), where RASP discriminates better in this case. Incorporating statistical potentials that weigh differently short, mid-range and long interactions like rsRNASP may not be the best choice for very close decoys.

Results are induced on the four scoring functions that we succeeded in implementing. We can not conclude on 3dR-NAscore, RNA3DCNN and ARES performances for ranking native-like structures.

#### H.3. Scoring functions and metrics relationship

We computed the ES and PCC scores for each data set, each available scoring function and metric. We considered for the metrics the RMSD, INF (also named INF_*all*_, as we considered the averaged value over base-pairing and base-stacking interactions), DI, MCQ, TM-score, GDT-TS, CAD-score, lDDT and *ϵ*RMSD. We did not include P-VALUE or CLASH score, as P-VALUE is like a condition-metric to assess the nonrandomness of the prediction. The CLASH score computation failed for most RNA molecules, leading to non-reliable results for this metric.

The results for the different datasets are given in Supplementary file. A summary of the most correlated metrics for each scoring function is provided in Table 4, where each best-related metric is counted for all three datasets. It shows that RASP has a high correlation with TM-score and MCQ in terms of ES, and MCQ, RMSD, lDDT and TM-score in terms of PCC. It means that RASP integrates atom deviation well in its statistical potential (as it is related to TM-score, RMSD, lDDT) and favours structures with good angle conservation (linked to MCQ). *ϵ*SCORE is linked to CAD-score, lDDT and TM-score in terms of ES, and INF_*all*_, MCQ and CAD-score. The high link with CAD-score in both evaluation criteria means it tends to conserve the RNA interactions (INF) and thus maintains a low contact-area difference (CAD-score). DFIRE-RNA is tied to CAD-score, lDDT and TM-score in terms of ES and MCQ, DI and lDDT for the PCC criteria. It seems to have some RNA interaction properties (INF/DI) while maintaining angle conservation (MCQ) and interatomic distance conservation (lDDT). Finally, rsRNASP is correlated to lDDT, TM-score in terms of ES, and INF_*all*_, *ϵ*RMSD, GDT-TS and CAD-score in terms of PCC. As rsRNASP considers low and high-range interactions, it tends to favour structures with good sequence alignment (lDDT, TM-score, *ϵ*RMSD, GDT-TS, CAD-score) and RNA structural features (INF).

**Table 4.**
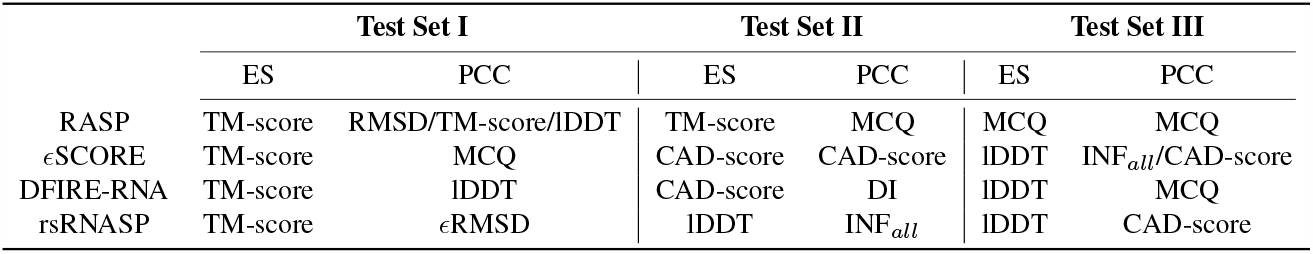
Summary of the best-correlated metrics for ES and PCC scores for each scoring function for the three test sets.

An example of RNA 1ec6D from Test Set I and its decoys is shown in Figure 3. We normalized all the scoring functions and computed the logarithm to plot them on the same scale. Growing scores were reversed to follow the same pattern as the others. DFIRE-RNA has a low slope compared to rsRNASP, RASP and *ϵ*SCORE. On the other hand, *ϵ*SCORE has a high slope and tends to increase the gap between near-native decoys. The overall high slope of *ϵ*SCORE for each metric shows a good discrimination property to divide native from non-native structures. It is supported by the number of native structures founded with the lowest *ϵ*SCORE for Test Set I: 85 out of 85.

**Figure 3.**
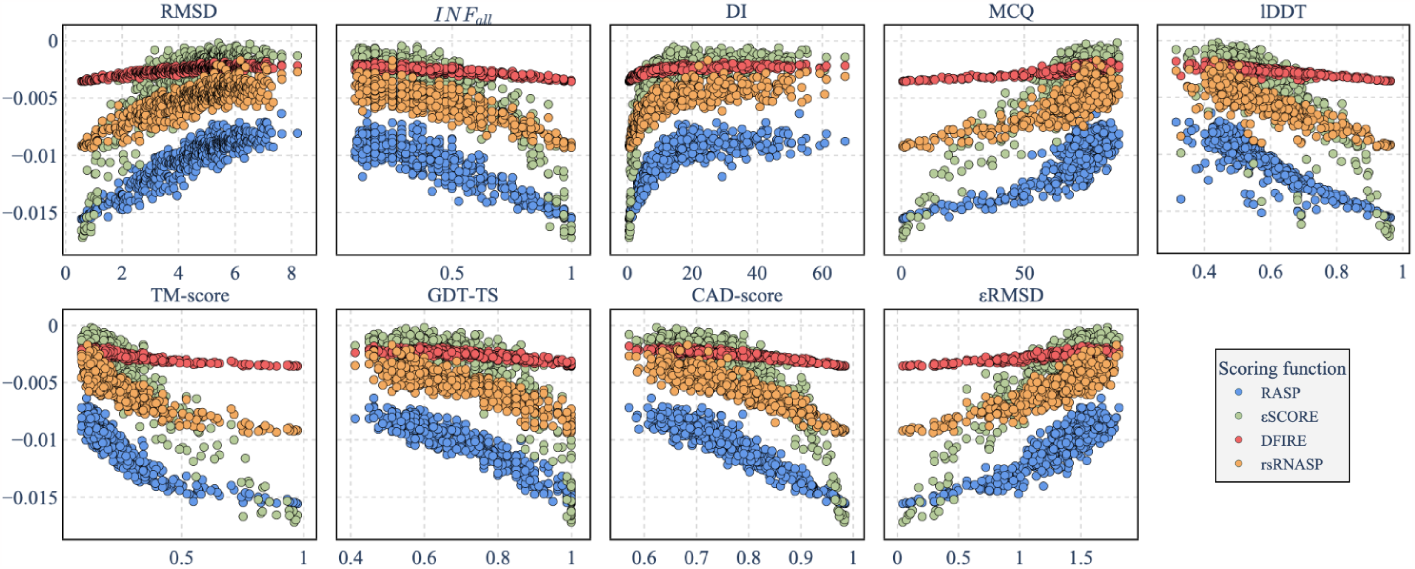
Logarithm of the normalized four scoring functions (RASP, *ϵ*SCORE, DFIRE-RNA and rsRNASP) of RNA 1ec6D and its 500 decoys from Test Set I depending on eight metrics. The increasing scores (like SCORE) were reversed to follow the same growth pattern as the other scoring functions.

#### H.4. Computation time and CO_2_ emissions

Computing a structure’s energy is integrated into developing models for predicting RNA 3D structures. Models for predicting 3D structures are usually slow and even slower when the sequence length increases. The computation time of energies shouldn’t be a bottleneck for selecting created models’ decoys.

We tracked the inference computation time for each energy for RNA of different lengths. We took as a benchmark the chain A with 2878 nucleotides of RNA 3f1hA from Test Set I. We randomly created five substructures for each step of 100 nucleotides from 100 to 2800. We tracked and averaged the time required to compute the scoring functions and metrics. It leads to Figure 4. It highlights the low computation time of DFIRE and *ϵ*SCORE that doesn’t exceed 20 seconds for RNA with a sequence length of less than 2800 nucleotides. The same goes for metrics like *ϵ*RMSD, GDT-TS, MCQ, DI, RMSD, and INF with a low computation time. On the other hand, RASP takes around 6min48 to compute for a sequence of 2800 nucleotides. rsRNASP has a complexity that almost explodes with the sequence length (10min39 for a sequence of 2800). This computation time is not scalable for the development of high-resolution models. For instance, if a predicted model generates 1000 structures of 2800 nucleotides and then tries to select the best ones with rsRNASP, it will take more than seven days and 9 hours. The MCQ and lDDT have a computation time higher than the other metrics for a sequence of more than a thousand nucleotides. MCQ has a computation time of less than 20 seconds compared to less than 1min17 for RNAs of 2800 nucleotides. CAD-score has a high computation time, with more than 2min20 for RNAs of 2800 nucleotides. Finally, the TM-score has a high computation time of 5min30 for a sequence of 2800 nucleotides.

**Figure 4.**
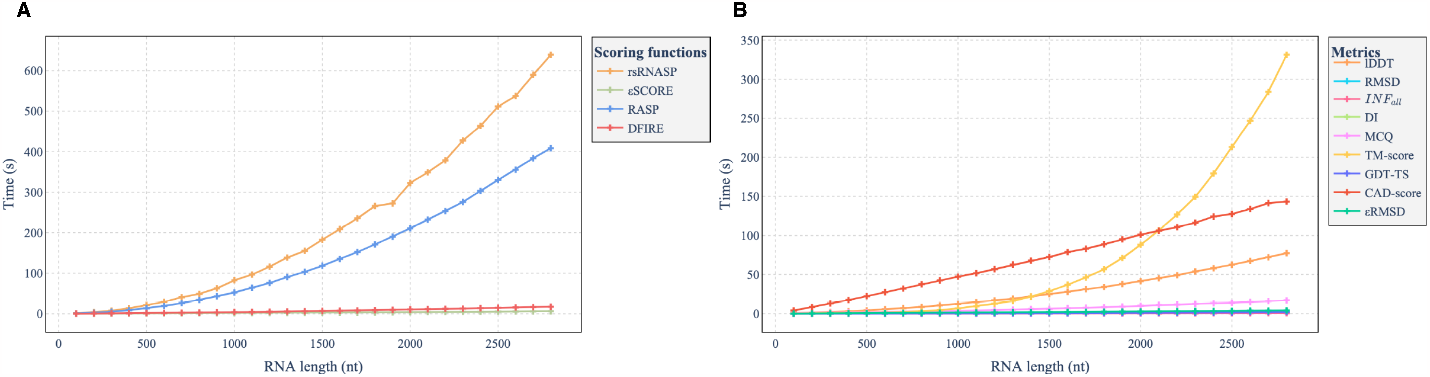
Computation time depending on the number of nucleotides in RNA sequences for substructures from RNA 3f1hA (Test Set I). A) Time executions for scoring functions. B) Time executions for metrics.

As computation methods can have environmental impacts, we also included carbon footprints of structural assessment methods. We computed for each dataset the equivalent CO_2_ measurements for an RNA (averaged over the different decoys), using CodeCarbon tool (42). The results are shown in Figure 5. We observe an overall higher consumption of CO_2_ for Test Set I, which is explained by the long RNAs in this dataset. rsRNASP and CAD-score have the highest CO_2_ consumption (with around 0.011 g/CO_2_ per RNA), followed by RASP and TM-score. CAD-score has a higher CO_2_ consumption compared to TM-score while calculating quicker. This difference could be explained by better resource management by the TM-score compared to the CAD-score.

**Figure 5.**
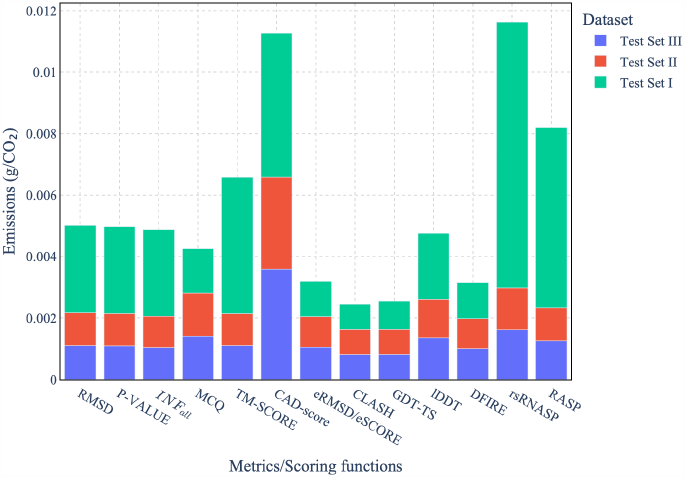
The CO_2_ equivalent measures for each metric and scoring function for each dataset. C*O*_2_ emissions

## I. Discussion

In these experiments, we found that rsRNASP outperforms the other scoring functions in finding the native structures among the three datasets while being correlated to lDDT, TM-score, INF, *ϵ*RMSD, GDT-TS and CAD-score. It comes with a price: its computation time highly increases with the number of nucleotides of an RNA sequence. Metrics also correlate between them: INF implies having good interaction accuracy, reducing the contact-area differences and the associated CAD metric. It is also correlated to *ϵ*RMSD, an improved RMSD considering the differences in the base interaction network. The MCQ metric seems to be the only metric that isn’t correlated to the others. Atomic deviation metrics tend also to be correlated: lDDT, TM-score, RMSD and *ϵ*RMSD.

We advise using MCQ as an evaluation metric to assume RNA nativity with DI (RMSD and INF) and lDDT (or TM-score or GDT-TS, as they are correlated). It provides a complementary set of metrics that assess RNA 3D structure evaluation compared to a reference structure. One should keep in mind the computation time and CO_2_ consumption that is associated with these metrics. Therefore, we do not recommend the CAD-score or TM-score, which have a high computation time and thus CO_2_ consumption.

As a scoring function, we suggest using rsRNASP. If the evaluated structures have a long sequence, we suggest using DFIRE-RNA or *ϵ*SCORE, which has good discriminating properties even if it doesn’t outperform rsRNASP. The consumption time and CO_2_ emissions of rsRNASP prevent using RNA with long sequences.

## Conclusion

In this work, we have presented a general overview of the assessment of the nativity of an RNA 3D structure. One can compare a predicted structure with comparative tools like atom distances, interaction accuracy or angle dissimilarity, given a reference structure. Such metrics can have general assumptions (like RMSD), whereas others tend to target RNA specificities. Protein metrics have also been adapted to be relevant for RNA assumption. Nonetheless, having a known solved structure is a strong condition and impossible when creating a model to predict RNA 3D structure. Instead, statistical potential energies tend to reproduce molecule-free energy: the lowest, the more stable and thus the more native a structure is. We have provided a review of the known RNA scoring functions and an extensive benchmark that is reproducible and open-source.

Each of these metrics and scoring functions results from years of research by different groups of researchers. Each code is written by different authors and is sometimes hard and time-consuming to install locally. We developed a software, named RNAdvisor, that gathers metrics and scoring functions in a unique interface. It provides a documented and wrapped code available in one command line. It helps centralize and automate the computation of metrics and scoring functions to assess RNA 3D structure nativity. RNAdvisor represents an advancement in the automation of RNA 3D structure evaluation. It facilitates the accessibility of existing metrics and scoring functions and thus can help accelerate investigation in RNA 3D structure predictions.

Future works could imply the development of new metrics that consider all the complementary specificities of RNA molecules and current metrics. This development must integrate existing metrics to avoid redundant work. The assessment of RNA nativity with energy score is still an area of research that should be explored. One should integrate the computation time to have a scoring function that could be adapted to long RNAs. Those developments should be guided with easy-to-use code to enable the reproducibility and integration of predicted models.

## ACKNOWLEDGEMENTS

This work is supported in part by UDOPIA-ANR-20-THIA-0013 and performed using HPC resources from GENCI/IDRIS (grant AD011014250). It was also partially supported by Labex DigiCosme (project ANR11LABEX0045DIGICOSME), operated by ANR as part of the program “Investissement d’Avenir” Idex ParisSaclay (ANR11IDEX000302).

## Footnotes

1 http://melolab.org/supmat/RNApot/Sup._Data.html.

2 https://github.com/Tan-group/rsRNASP.

3 https://github.com/RNA-Puzzles/standardized_dataset/tree/master

