## Supplementary materials for "RNAdvisor: a comprehensive benchmarking tool for the measure and prediction of RNA structural model quality"

### Supplementary Note 1: State-of-the-art metrics

#### A. General metrics

***A.1. Root-Mean-Square Devation (RMSD).*** The RMSD is described as the Root-mean-square deviation of two molecules. It computes the mean Euclidean distances of the aligned atoms after superimposition. The alignment centers and rotates the two molecules using the Kabsch algorithm (1). The following formula defines it:

$$RMSD(X,Y) = \sqrt{\frac{\sum_{i=1}^{N}(X_i - Y_i)}{N}}$$

where $X$ is the predicted structure, $Y$ the reference and $X_i$ the $i-th$ atoms of the molecule. Note that X and Y should be aligned, otherwise, the RMSD is computed with a rotation matrix added to it:

$$RMSD(X,Y) = \min(\sqrt{\frac{\sum_{i=1}^{N}(X_i - RY_i)}{N}})$$

with R a rotation matrix.

RMSD ranges from 0Å to any positive values. As crystallographic molecules have an approximation of about 2Å, an RMSD around this value means a confident prediction. This metric is the most used across the community, giving a quick idea of how good the prediction is. On the other hand, it doesn't say how 'bad' the prediction could be: a high RMSD doesn't indicate the failing components. A molecule can have a high RMSD while only having a small portion of the structure that differs from the native one.

***A.2. $\epsilon$RMSD .*** $\epsilon$RMSD (2) is an improved RMSD that tries to solve one of the main issues of RMSD: the lack of reliable information about the differences in the base interaction network. It uses a specific molecular representation of the bases, with the local coordinate system in the center of six-membered rings. The relative orientation and positions between nucleobases can thus be defined by a vector $r$, in cylindrical coordinates $\rho$, $\theta$ and $z$. This coordinate system defines almost all base-stacking and base-pairing interactions in a well-defined ellipsoidal shell. The anisotropic position vector is then defined as:

$$\tilde{r} = (\frac{r_x}{a}, \frac{r_y}{a}, \frac{r_z}{b}) = (\frac{\rho}{a}\cos\theta, \frac{\rho}{a}\sin\theta, \frac{z}{b})$$

This provides information about the relative base arrangement, enabling the definition of a RMSD-like metrics:

$$\epsilon\text{RMSD}(X,Y) = \sqrt{\frac{1}{N}\sum_{i,j}|G(\tilde{r}_{ij}^{X}) - G(\tilde{r}_{ij}^{Y})|^2}$$

with $\tilde{r}_{ij}^{X}$ is the $i$th base of structure X paired with the $j$th pair of reference structure Y. $G$ is a function that tries to avoid the significant deviation in distant pairs. Using a $\tilde{r}_{cutoff}$ value for the distance would bring discontinuity in the metric function, that is why they ended up with the $G$ function defined by:

$$G(\tilde{r}) = \begin{pmatrix} sin(\gamma\tilde{r})\frac{\tilde{r}_x}{\tilde{r}} \\ sin(\gamma\tilde{r})\frac{\tilde{r}_y}{\tilde{r}} \\ sin(\gamma\tilde{r})\frac{\tilde{r}_z}{\tilde{r}} \\ 1 + cos(\gamma\tilde{r}) \end{pmatrix} \times \frac{\Theta(\tilde{r}_{cutoff} - \tilde{r})}{\gamma}$$

with $\gamma = \frac{\pi}{\tilde{r}_{cutoff}}$ and $r_{cutoff}$ a constant equal to 2.4 after analysis. $\Theta$ is the Heaviside function.

This metric considers nucleobases' relative distance and orientation because of the $\tilde{r}$ distance coordinate. It is close to the INF score discussed above while being a continuous function of the atomic coordinates.

***A.3. CLASH.*** The clashscore (3) is an overall metric of the number of overlaps >0.4Å per thousand atoms. A steric clash is an unnatural overlap of two non-bonding atoms in a structure. It can exist in any structure predicted using computation methods and even in experimental data like NMR or X-ray. The clashscore was thus developed to identify non-native atom positions.
It has been developed with MolProbity (3), a web-server

that offers quality validation for 3D structures of proteins, nucleic acids and complexes.

### B. Protein-inspired metrics

***B.1. Template Modeling score (TM-score).*** The TM-score (4) is a widely used metric in the CASP competition. Instead of using the RMSD that doesn't consider the residual alignment, the TM-score considers both residual alignment coverage and distance normalization. As the normalisation parameters are specific to proteins, an adaptation of the TM-score was introduced with RNA-Align (5), a work that produces RNA alignments. The given formula defines TM-score for RNA:

$$\text{TM-score}_{RNA} = \frac{1}{L}\sum_{i=1}^{L_{ali}} \frac{1}{1+(\frac{d_i}{d_0})^2}$$

with $L$ the length of the target RNA, $L_{ali}$ the number of aligned nucleotides, and $d_i$ the distance between the $i-th$ aligned pair of residues. The scaling factor $d_0$ prevents the score from being dependent on the length of the RNA. It is defined as :

$$d_0 = 0.6\sqrt{L-0.5} - 2.5$$

The parameters were derived from a large PDB set of random RNA pairs. The alignment used is a heuristic dynamic programming iteration process assisted by distance-based secondary structure assignments.

***B.2. Global Distance Test Total Score (GDT_TS).*** The Global Distance Test Total Score is derived from a method of alignment of longest continuous sequences: LGA(Local-Global Alignment) (6). The Global Distance Test (GDT) is a score that estimates the percent of residues that can fit under a distance cutoff using different superimpositions. The GDT_TS is then computed as the average over the threshold of 1, 2, 4 and 8 Å:

$$GDT\_TS = \frac{P_1 + P_2 + P_4 + P_8}{4}$$

where $P_d$ is the percent of residues from a candidate that can be superimposed with corresponding residues in the target structure under a distance cutoff of $d$ Å. It ranges from 0 (bad prediction) to 1 (perfect prediction). Random predictions tend to give a GDT_TS score of around 0.2, while getting the rough topology of the molecule usually gives a score around 0.5. This metric is widely used in the CASP competition. This score gives an overall idea of how well a prediction can superimpose a reference. Nonetheless, it doesn't explicit the regions with good matching as it is averaged over different cutoffs.

***B.3. Contact-area Difference score (CAD-score).*** The CAD-score (7, 8) measures the structural similarity in a contact-area difference-based function. This metric is based on the work of (9), which introduced a residue-residue contact area score to compare a structure to a reference. The CAD-score is defined with the set of all pairs of residues $(i,j)$, denoting G, that have non-contact area $T_{(i,j)}$ in the target structure. The contact area of the residue pairs $(i,j) \in G$ is described as $M_{(i,j)}$ for the candidate structure. The contact area difference (CAD) is thus defined as the absolute difference of contact areas between the residues $(i,j)$ in target T and candidate structure M:

$$CAD_{(i,j)} = |T_{(i,j)} - M_{(i,j)}|$$

A bounded value is used to prevent over and under-prediction of contact areas:

$$CAD_{(i,j)}^{bonded} = \min(CAD_{(i,j)}, T_{(i,j)})$$

Then, the final CAD-score is computed as:

$$CAD - score = 1 - \frac{\sum_{(i,j)\in G} CAD_{(i,j)}^{bonded}}{\sum_{(i,j)\in G} T_{(i,j)}}$$

The CAD-score ranges between 0 and 1, where 1 means the prediction and the target are identical. The computation process of CAD-score is directly adaptable to RNA molecules, which has been done with the web server (8). One of the main advantages of the CAD-score is the display of a stronger emphasis on the physical realism of models.

***B.4. Local Distance Difference Test.*** The local distance difference test (lDDT) (10) assesses the interatomic distance differences between a reference structure and a predicted one. It does not require any superposition. The lDDT considers all the pairs of atoms in the reference structure within a $R_0$ distance, where $R_0$ (the inclusion radius, default value of 15 Å) is a predefined threshold. The atom pairs define a set of distances $L$, which is used for a predicted model. A distance in the prediction is preserved if, given a threshold, it is the same as its corresponding distance in $L$. The lDDT is thus derived using four different thresholds: 0.5 Å, 1 Å, 2 Å, and 4 Å. lDDT is the average of four fractions of conserved distances within the four thresholds. It ranges between 0 and 1, where 1 means a perfect reconstruction of interatomic distances.
lDDT captures local atomic interactions and is robust to outliers. One of the main advantages is that it quantifies the model quality on the level of the residue's environment.

### C. RNA-oriented metrics

***C.1. P-value.*** The P-value (11) assesses the probability that a given structure is better than that expected by chance. It is based on an empirical law relationship between mean RMSD and chain length. The original paper finds its law by generating decoy structures using replica exchange discrete molecular dynamics simulation. They then used the mean and standard deviation distribution of each RMSD and derived an expression that relates the RMSD to chain length. The relation they found is the following:

$$\langle RMSD \rangle = aN^{(0.41)} - b$$

where $N$ is the length of the chain, $a$ and $b$ are constants that depend on the provided secondary structure as inputs to the molecular dynamic simulation. The P-value is then computed as the RNA prediction significance from the Z-score, given a predicted structure that differs from an accepted structure by an RMSD of m:

$$P-value = \frac{1+erf(\frac{Z}{\sqrt{2}})}{2}$$

where

$$Z = \frac{m - \langle RMSD \rangle}{\sigma_m}$$

with $\sigma_m \approx 1.8$Å.

P-value expresses the non-randomness of RNA structure prediction. A P-value below 0.01 represents a successful prediction for RNAs between 35 and 161 nucleotides.

***C.2. Interaction Network Fidelity (INF).*** The Interaction Network Fidelity (INF) (12) is a metric that considers the specific RNA structural features such as helices or hairpins. It considers two main interactions for RNA molecules: base-stacking and base-pairing. Using Leontis and Westhof nomenclature (**?** ), each base interacts with one of three edges: Watson-Crick edge, Hoogsten edge, and sugar edge. Each type of edge has an orientation relative to the backbone: either *cis* or *trans*. Given the two orientations and the three edges of each base, there are 12 possible base pairs. The classic base pairing is defined as the Watson-Crick cis pairing with another base, whereas the others are non-canonical. The INF metric integrates these types of interactions in the scoring process.

If we denote $S_m$ the different interactions of the candidate structure, and $S_r$ those of the reference solved structure, we can define the interactions of both sets as $TP = S_r \cap S_m$. It defines the true positives, while the false positives are expressed as $FP = S_m \backslash S_r$. The interactions present in $S_r$ but not in $S_m$ are the false negatives: $FN = S_r \backslash S_m$. The INF score is then defined as:

$$INF(A,B) = MCC(A,B)$$

with $MCC$ the Matthews Correlation Coefficient:

$$MCC = \sqrt{PPV \times STY}$$

The PPV denotes the specificity:

$$PPV = \frac{|TP|}{|TP| + |FP|}$$

and the STY defines the sensitivity:

$$STY = \frac{|TP|}{|TP| + |FN|}$$

When the predicted model reproduces exactly the reference interactions, $|TP| > 1$ and $FP = FN = 0$. The $MCC$ value equals 1, such as the INF score. On the other hand, if none of the interactions of the reference is reproduced in the predicted model, $|TP| = 0$, and the INF score equals 0. The INF score can be specific to base-pairing interactions (INF$_{bp}$), the base stacking interactions (INF$_{stack}$), or consider both (INF$_{all}$).

***C.3. Deformation Index (DI).*** INF score alone is not enough to assess RNA 3D structure quality. RMSD, on the other hand, can't consider the local dissimilarity as it averages the error through the entire structure. The Deformation Index (DI) aims to take the best of those two metrics. It is defined as:

$$DI(A,B) = \frac{RMSD(A,B)}{INF(A,B)}$$

Nevertheless, the aim of having a unique value to assess structure dissimilarities is limited. Local or global conformation shape is complex, and each domain has specificities. The automation of dissimilarity assessment can't be made if we rely on human intervention to assess and understand each prediction's mistakes. The deformation index is, therefore, a good tradeoff to have a quick overview of the quality of a prediction. It encodes the overall quality of the prediction divided by the quality of the reproduced interactions.

***C.4. Deformation Profile (DP).*** The deformation profile (DP) considers dissimilarities between structures at the nucleotide scale, considering interdomain and intradomain interactions. This matrix highlights the average distance between a predicted model (PM) and the reference model (RM). Given the $i$th nucleotide from the predicted model, noted $PM_i$, and the nucleotide from the reference model, $RM_i$, the model that superimposes $PM_i$ over the $RM_i$ is noted as $SUP(RM_i, PM_i)$. This superimposition is executed by minimizing the RMSDs between all the atoms of the nucleotides between the two models. The DP is then computed as:

$$DP_{i,j} = \text{AVG\_DIST}[\text{SUP}(RM_i, PM_i)_j, RM_j]$$

where AVG_DIST is the average distance between all atoms of the nucleotides of the two models. The $i$th row of the matrix gives an estimation of the local similarity around the $i$th nucleotide, where this nucleotide is superimposed with the reference $i$th nucleotide. The $j$th column gives the average atomic distance between the $j$th nucleotides of both the reference, the prediction for each superimposition. The diagonal gives insights into individual nucleotide conformation similarity. Nevertheless, the DP is not normalized, meaning the values near the diagonal tend to be smaller than those farther away. This is due to the atomic distance computed after superimposed: closed nucleotides tend to be closer than those far away from this nucleotide.

***C.5. Mean of Circular Quantities (MCQ).*** The mean of circular quantities (MCQ) is a metric that uses algebraic and trigonometric space for the dissimilarity measure. It considers a molecule 3D structure as a set of torsional angles. Each RNA residue is described by eight torsion angles: the seven classic RNA torsion angles ($\alpha$, $\beta$, $\gamma$, $\delta$, $\epsilon$, $\xi$ and $\chi$) and the sugar pucker pseudorotation phase called $P$. The $P$ torsion angle is used to define the ribose ring. To compute similarity between trigonometric structure $S_T$ and $S_{T'}$, it considers $2\pi$-periodical circular quantities. To compare two circular quantities $t$ and $t'$, the difference is defined as:

$$\text{diff}(t,t') = |\text{mod}(t) - \text{mod}(t')|$$

with:

$$\text{mod}(t) = (t + 2\pi) \quad \text{modulo } 2\pi$$

Therefore, the distance between two angles $t$ and $t'$ is described as:

$$\Delta(t,t') = \begin{cases} 0 & \text{if t, t' undefined} \\ \pi & \text{if t or t' undefined} \\ \min(\text{diff}(t,t'), 2\pi - \text{diff}(t,t')) & \text{otherwise} \end{cases}$$

The sum of differences of angles, $D_{sin}$ and $D_{cos}$, defined by:

$$D_{sin} = \frac{1}{r|T|} \sum_{i=1}^{r} \sum_{j=1}^{|T|} \sin \Delta(t_{i,j}, t'_{i,j})$$

$$D_{cos} = \frac{1}{r|T|} \sum_{i=1}^{r} \sum_{j=1}^{|T|} \cos \Delta(t_{i,j}, t'_{i,j})$$

Finally, the overall distance between structures $S_T$ and $S_{T'}$ is given by:

$$MCQ(S_T, S_{T'}) = \arctan(D_{sin}, D_{cos})$$

with $r$ the number of residues in $S \cap S'$ and $T$ the set of torsion angles. This metric is computed using the MCQ4Structures algorithm. The MCQ gives a global dissimilarity measure. One of the advantages of the MCQ score is the possibility to consider inputs of different forms. Indeed, as it uses trigonometric space, it is possible to have different types of representation for a 3D structure and thus compute the score. The other benefit is that it complements other metrics, giving higher scores for structures with common shapes. It doesn't require superimposition, leading to less computation.

### Supplementary Note 2: State-of-the-art scoring functions

#### A. Knowledge-based scoring functions

***A.1. Ribonucleic Acids Statistical Potential (RASP).*** RASP (13) is an all-atom knowledge-based potential. It uses a pairwise distance-dependent energy score. The dataset employed was decoys created from 85 native structures from the PDB. Decoys were generated using Gaussian restraints for dihedral angles and atom distances. The distance considered for a given pair of atoms is discretized in the 0 to 20 Å range, with an interval of 1Å. They used the average method to get a reference state. Parameters of the energy score have been optimized with the same set of native structures used to derive the potentials. The critical parameters of the energy score are the distance-dependent descriptors of atom pairs, the sequence separation to account for local or non-local interactions, and the atom types.

***A.2. $\epsilon$SCORE.*** The $\epsilon$SCORE was introduced with the $\epsilon$RMSD metric (2). It is based on the six-membered rings coordinate system, where each relative orientation between two nucleobases is described with the $r$ vector. They ended up with a knowledge-based scoring function that accounts for the compatibility of an RNA molecule towards the expected distribution observed on a set of native structures. It is defined as:

$$\epsilon\text{SCORE} = \sum_{j,k} p(r_{j,k})$$

with $p(r)$ the empirical probability distribution of nucleobases in the crystal structure of H. The sum is used instead of the product to reduce the effect of low-count regions. This scoring function uses a minimalist description of the structure to get a score that has proven to be efficient.

***A.3. 3dRNAScore.*** 3dRNAScore (14) is a knowledge-based score that uses distance-dependent and dihedral-dependent energies. The first part of the energy uses the distance between any non-bonded atom in different residues through the molecule. It is based on the Boltzmann distribution. Indeed, one assumption is that the relative free energy of a structure can be deduced from the inverse of Boltzmann's law. Given a distance $d$ between atoms $i$ and $j$ from residue $a$ and $b$, the free energy can be approximated by:

$$\Delta G(d) = -k_B T \sum_{i,j} \ln \frac{f_{ab}^{OBS}(d_{ab}^{ij})}{f_{ab}^{REF}(d_{ab}^{ij})}$$

where $T$ is the constant temperature, set as $298K$ and $k_B$ the Boltzmann's constant. $f_{ab}^{OBS}(d_{ab}^{ij})$ represents the probability of the distance $d^{ij}$ between atoms $i$ and $j$ of type $a$ and $b$ in the native RNA structures. On the other hand, $f_{ab}^{REF}(d_{ab}^{ij})$ represents the same distance but for an RNA structure that is not native, named the reference state. 3dRNAScore used the average reference state, meaning that the atom types are ignored. 3DRNAScore considers the atom pair in two adjacent bases, unlike other statistical potentials. The distance cutoff is 20 Å with a bin width of 0.15 Å.

The other term of the energy stands for the dihedral potential. Is uses the seven torsion angles of RNA ($\alpha$, $\beta$, $\gamma$, $\delta$, $\epsilon$, $\xi$ and $\chi$). They assumed that these angles follow the Boltzmann statistical distribution:

$$\Delta G_i(\theta_a^i) = -k_B T \ln \frac{f_a^{OBS}(\theta_a^i)}{f_a^{REF}(\theta_a^i)}$$

where $f_a^{OBS}(\theta_a^i)$ is the observed probability of angle $i$ of $\theta$ degree (0-360) and type $a$ in the database of RNA structures. The bin width used is 4.5°.

The final formula for the 3dRNAScore is a weighted sum of the two previous mean force potentials:

$$\Delta G_{\text{total}} = \Delta G_{\text{distance}} + \omega \Delta G_{\text{dihedral}}$$

where $\omega$ is a parameter optimized to maximize the correlation between 3dRNAScore and the DI metric. They used a non-redundant set of 317 RNA structures derived from the PDB to compute their score. One of the advantages of 3dRNAScore is the account of both base pair and dihedral angles. The availability of RNA structures nonetheless limits knowledge-based energies.

The code source seems available, but we didn't manage to download it and extract it. It requires having an account. When logged, the available source code seems not downloadable, as the compressed folder can't be extracted (it is a 0-byte content).

***A.4. DFIRE-RNA.*** DFIRE-RNA (15) is an all-atom, distance-dependent, knowledge-based energy function. They considered that the reference state used by 3dRNAScore and RASP, the average state, is not optimal as the average interaction is not always zero. They developed a distance-scaled, finite ideal-gas reference as the reference state should be without interactions. The DFIRE-RNA energy score is defined as follows:

$$F(r_{ij}|a_i,a_j) = -RT \ln \frac{N_{obs}(r_{ij}|a_i,a_j)}{(\frac{r_{ij}}{r_{cut}})^{\alpha} \frac{\Delta r}{\Delta r_{cut}} N_{obs}(r_{cut}|a_i,a_j)}$$

with $N_{obs}(r_{ij}|a_i,a_j)$ is the observed number of atomic pairs ($a_i$, $a_j$) within a distance $r_{ij}$. They used all 85 atom types, where each atom in a nucleotide is considered different. The bin width $\Delta r$ is equal to 0.7Å. The distance cutoff is set to 19Å, and the $\alpha$ is set to 1.61. They used a dataset of 405 non-redundant RNA molecules derived from the PDB.

One limitation of DFIRE-RNA is the dependence on distances. Nonetheless, RNA structures are mainly based on hydrogen-bonded base pairing and stacking, which depend more on orientation. The distance-dependent can't fully consider the basement of RNA folding stabilization.

***A.5. rsRNASP.*** rsRNASP (16) is an all-atom pairwise-dependent knowledge base score. It states that no statistical potential considers the range of residue interactions. Thus, it integrates a separation between short, medium and long-range interactions into the scoring function. A separation threshold $k_0$ was used to classify an interaction as short- or long-range. The energy for a conformation C is given by:

$$\Delta E(S) = \sum_{k>2}^{k \leq k_0} \Delta E_{short}(i,j,r) + w \sum_{k>k_0} \Delta E_{long}(i,j,r)$$

where:

$$\Delta E_{short}(i,j,r) = -k_B T \ln \frac{P^{OBS}_{2<k\leq k_0}(i,j,r)}{P^{REF}_{2<k\leq k_0}(i,j,r)}$$

and

$$\Delta E_{long}(i,j,r) = -k_B T \ln \frac{P^{OBS}_{k>k_0}(i,j,r)}{P^{REF}_{k>k_0}(i,j,r)}$$

with $w$ the weight to balance the contribution of the long-range interactions.

They used two reference states: the random-walk-chain (17) and averaging (18) to build the long-ranged and short-ranged potentials. They derived 191 non-redundant RNA structures as a training set from the PDB. The distance bin width was set to 0.3Å, and the distance cutoffs were set to 13Å and 24Å for short- and long-range interactions, respectively. The key advantages of rsRNASP are separating short- and long-range interactions, such as the reference states used. The limitations are the lack of geometrical parameters in the energy, like torsion angles. Those angles play a crucial role in determining the RNA 3D motifs. Finally, the last limit is the number of available RNAs.

### B. Deep learning scoring functions

***B.1. RNA3DCNN.*** The different advances inspire RNA3DCNN in terms of deep learning. They assume that an RNA molecule can be seen as a 3D image, and thus convolutional neural networks could help infer information. The method relies on the fact that each RNA molecule has a different global shape and similar local conformation. The RNA3DCNN uses as inputs a cube of local atoms and outputs an RMSD-based residue unfitness score. The convolutional model outputs an unfitness score for each local cube of atoms that are then summed up to give a global score to the structure. A Cartesian coordinate centered at the $C_1'$ atom is implemented, where all the atoms of a distance below 16Å are considered. They create a grid of 32x32x32 Å, each comprising voxels of three channels: occupation number, mass and charge of the present atoms. The channels are inspired by the RGB channels used in images. The model uses four 3D convolutional layers, with filters of size 8, 16, 32 and 64. The output layer returns a unique value named the unfitness score. 332 non-redundant RNA molecules were extracted from the PDB, while 82 RNAs were used as validation sets. They augmented the available data with molecular dynamics simulation for each native data and kept the best structures based on RMSD. They also used a Monte Carlo process to generate other decoys. They trained their RNA3DCNN score separately for each augmentation dataset, leading to RNA3DCNN_MD and RNA3DCNN_MDMC. The main advantage of this scoring function is that it doesn't require a reference state, compared to traditional statistical potentials. It can also evaluate each nucleotide, meaning that it could guide the refinement of structure.

***B.2. Atomic Rotationally Equivariant Scorer (ARES).*** The Atomic Rotationally Equivariant Scorer (ARES) (19) is a scoring function incorporating only atomic coordinates and chemical element type as inputs. It is trained with 18 RNA structures, augmented by 1000 decoys for each structure with FARFAR 2 (20). They used the equivariance property of specific neural networks, where rotation or translation in the 3D space (and network inputs) results in the same transformation in the output. The first layer of the network collects information about local properties (like position and orientation), while the latter layers infer global information. The network's output is the RMSD, while the inputs use coordinates and chemical properties. It is passed through an equivariant convolution (from Tensor Field Network) that learns features to close atoms. A radial function is added to consider the distance relationship between atoms (but not the orientation). It is associated with an angular function that includes only orientation between atoms. One of the advantages of ARES is the inferred representations that are coming from deep neural networks. While other scores require the account of specific interactions, ARES lets the neural networks find them. The drawback could be the non-understanding of all the features used. It could also be biased by the database used: decoys from a unique method, FARFAR 2.

### Supplementary Note 3: RNAdvisor tool

#### A. Code integration

RNAdvisor is a Docker-based tool that gathers nine source codes written in Java, C++ and Python into one interface. We modified some codes to integrate them properly into the common interface.

We describe in the following the different codes used for the computation of metrics and energies.

**RNA_assessment**: A Python-based repository that is almost complete for the computation of metrics. We did some modifications to make the code more readable (by creating files that separate the classes). The modified code is available in a Github fork. It computes the CLASH, P-VALUE, RMSD, INF and DI scores. It is also used to normalize the structures, which is integrated and added for all the following code.

**BaRNAba**: A Python-based repository that computes the $\epsilon$RMSD and $\epsilon$SCORE. We created a fork to remove the non-wanted prints and change the signature of some functions to integrate it well. A Github fork is available.

**MCQ4Structures** : A Java-based code that computes the MCQ score. We executed the code using the command line and thus didn't require any modification. We parsed the output to get the metric.

**Voronota**: A C++ code that computes the CAD-score. We didn't have to modify the source code as we executed it with the command line. We parsed the output using *awk* command.

**ZhangLab**: A C++ script that comes from a web server. Multiple scoring functions are available on this website but are adapted to proteins. This script computes the GDT-TS and is also adapted to RNA.

**OpenStructure**: A C++ and Python program that computes multiple protein-based scoring functions. We integrated the Docker image as the based image. It computes the lDDT and TM-score.

**RASP**: A C++ code that computes the RASP energy. It comes from a web server. It requires a specific version of g++ compiler. Therefore, We created a Github fork. We added all the installation processes that I summarized with a Makefile and a Dockerfile.

**rsRNASP**: A C++ code to compute the rsRNASP energy. It provides the different datasets used for training and evaluation. We removed the non-needed *pdb* files that could increase the total size of the Docker image. The reduced code is available in a Github fork.

**DFIRE-RNA**: A C++ code that implements the DFIRE energy. We didn't modify the code and only parsed the output.

#### B. Usage

For the metrics computation, it takes as inputs two *pdb* files (the reference and the prediction) and returns a *csv* with the different metrics. The energy computation takes one *pdb* file as input. By default, when two files are provided as inputs, it will also compute the energies. To provide an easy way to compute these metrics and energies for multiple predictions, it is also possible (and recommended) to provide as inputs a *pdb* file (for reference) and a path to a folder with multiple predictions. Table 1 describes the different command line arguments.

| Arguments | Description |
|---|---|
| pred_path | Directory to *pdb* files |
| native_path | Path to the native structure |
| result_path | Path to store the different scores |
| time_path | Path to store the time of each metric |
| log_path | Path to store the log of the script |
| all_scores | List of the scores to use |
| no_normalisation | Normalisation of the files |
| sort_by | Metric to sort the results by |
| config_path | Path to the *config.yaml* file |
| verbose | Whether to print the debug logs |

**Table 1.** Arguments of the RNAdvisor code in command line

The user has multiple options to call the program. It can use direct Python commands with different arguments. The user can also provide all the parameters in a *config.yaml* file. The user can also execute the code using *docker run* command. It should thus provide a path to the local files not part of the Docker image.

### Supplementary Note 4: Results

#### A. Metrics relationship

Figure 1 shows the ES and PCC between each metric for each dataset. We observe a similar behaviour between Test Set II and Test Set III for both ES and PCC compared to the ones from Test Set I. Indeed, the MCQ correlation score is lower for Test Set II and III, which is not the case for Test Set I. This difference can be explained by the decoys used for Test Set II and III: structures from predictive models whereas the decoys from Test Set I are more near-native structures. In terms of MCQ, as predictions from Test Set II and III are not derived from the native ones, the angle conservation is poor, and thus, the metric hardly discriminates the different structures.

### B. Scoring function ranking

The average native rank for each dataset is shown in Table 2. It shows that RASP struggles for Test Set III, with an average native rank of 16.41, whereas it has a good performance for Test Set I. rsRNASP has a superior ranking performance with an average native rank of 3.64 for Test Set III. $\epsilon$SCORE and DFIRE have slightly similar rank for native structure, with a slightly better average position for $\epsilon$SCORE on Test Set II and III.

| | Energies | | | |
|---|---|---|---|---|
| Dataset | RASP | $\epsilon$SCORE | DFIRE | rsRNASP |
| Test Set I | 1.18 | **1** | 1.04 | 1.48 |
| Test Set II | 11.2 | 5.1 | 4.15 | **1.95** |
| Test Set III | 16.41 | 7 | 7.82 | **3.64** |

**Table 2.** Average energy position of the native structures found for each dataset.

***B.1. Scoring functions and metrics relationship.*** For Test Set I, the TM-score has the best correlation, in terms of ES, for all four scoring functions. It means that the top 10% of structures from the scoring functions intersect best with the top structures ranked by the TM-score. The correlation of all ranking structures is different depending on scoring functions: RASP correlates best to RMSD, TM-score and lDDT (PCC of 0.9). $\epsilon$SCORE is best related MCQ (PCC of 0.91), whereas DFIRE-RNA correlates to lDDT. Finally, rsRNASP is tied with $\epsilon$RMSD (PCC of 0.87), whereas it is also close to MCQ, TM-score and lDDT (PCC of 0.86).

For Test Set II, the distribution of correlation differs for each scoring function. The overall values of ES and PCC for this dataset are significantly lower than for Test Set I, as the decoys are generated differently. Test Set I comprises near-native structures with small perturbations compared to Test Set II and Test Set III, which have structures from different predictive models. RASP seems more correlated to MCQ (PCC of 0.48) and TM-score (ES of 2.36), while $\epsilon$SCORE has a higher correlation with CAD-score (ES of 2.37 and PCC of 0.38). DFIRE-RNA is related to CAD-score (ES of 4.09) and DI (PCC of 0.55), while rsRNASP is connected to lDDT (ES of 5.1) and $INF_{all}$ (PCC of 0.66).

For Test Set III, RASP is correlated to MCQ (ES of 2.36 and PCC of 0.65) and $\epsilon$SCORE to $INF_{all}$, CAD-score (PCC of 0.53) and lDDT (ES of 3.44). DFIRE-RNA is linked to lDDT (ES of 4.39) and MCQ (PCC of 0.58), and rsNASP to CAD-score (PCC of 0.67) and lDDT (ES of 5.29).

The results for the different datasets are given in Table 3.

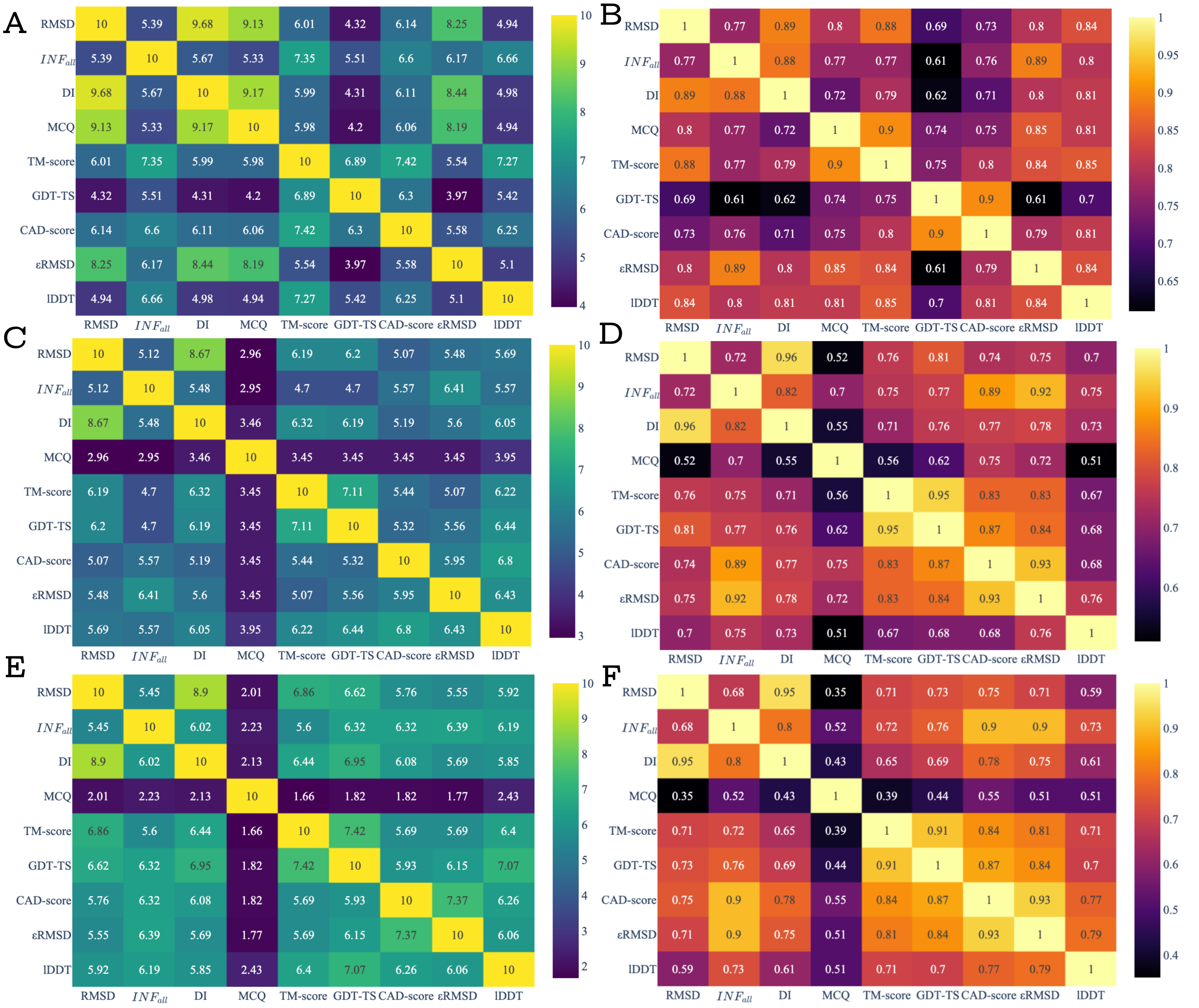


**Figure 1.** PCC and ES between each metric for each dataset. A) and B): ES results, respectively PCC results, for Test Set I. C) and D): ES results, respectovely results, for Test Set II. E) and F): ES results, respecively PCC for Test Set III.

| | RMSD | | $INF_{all}$ | | DI | | MCQ | | TM-score | | GDT-TS | | CAD-score | | ϵRMSD | | lDDT | |
|---|---|---|---|---|---|---|---|---|---|---|---|---|---|---|---|---|---|---|
| **Test Set I** | | | | | | | | | | | | | | | | | | |
| | ES | PCC | ES | PCC | ES | PCC | ES | PCC | ES | PCC | ES | PCC | ES | PCC | ES | PCC | ES | PCC |
| RASP | 5.95 | **0.9** | 7.77 | 0.84 | 6.05 | 0.86 | 6.11 | 0.88 | **8.73** | **0.9** | 6.49 | 0.71 | 7.69 | 0.85 | 5.88 | 0.89 | 7.45 | **0.9** |
| ϵSCORE | 5.75 | 0.82 | 7.81 | 0.82 | 5.88 | 0.77 | 5.91 | **0.91** | **8.43** | 0.89 | 6.25 | 0.66 | 7.49 | 0.81 | 5.9 | 0.9 | 7.29 | 0.85 |
| DFIRE-RNA | 5.71 | 0.84 | 7.51 | 0.82 | 5.81 | 0.81 | 5.91 | 0.87 | **8.28** | 0.86 | 6.22 | 0.68 | 7.32 | 0.83 | 5.73 | 0.87 | 7.15 | **0.88** |
| rsRNASP | 5.78 | 0.84 | 7.68 | 0.81 | 5.89 | 0.81 | 5.89 | 0.86 | **8.37** | 0.86 | 6.24 | 0.68 | 7.39 | 0.82 | 5.84 | **0.87** | 7.21 | 0.86 |
| **Test Set II** | | | | | | | | | | | | | | | | | | |
| | ES | PCC | ES | PCC | ES | PCC | ES | PCC | ES | PCC | ES | PCC | ES | PCC | ES | PCC | ES | PCC |
| RASP | 1.73 | 0.32 | 1.84 | 0.37 | 1.73 | 0.38 | 2.23 | **0.48** | **2.36** | 0.25 | 1.97 | 0.26 | 1.97 | 0.36 | 1.97 | 0.3 | 2.12 | 0.23 |
| ϵSCORE | 1.62 | 0.3 | 2.12 | 0.36 | 1.75 | 0.33 | 1.12 | 0.37 | 2.11 | 0.33 | 2.12 | 0.34 | **2.37** | **0.38** | 1.99 | 0.28 | 2.24 | 0.19 |
| DFIRE-RNA | 3.1 | 0.54 | 2.98 | 0.52 | 3.23 | **0.55** | 1.97 | 0.53 | 3.86 | 0.5 | 3.47 | 0.52 | **4.09** | 0.52 | 3.23 | 0.46 | 4 | 0.52 |
| rsRNASP | 4.09 | 0.63 | 4.09 | **0.66** | 4.09 | 0.64 | 2.71 | 0.6 | 4.61 | 0.63 | 4.6 | 0.66 | 4.59 | 0.65 | 4.22 | 0.61 | **5.1** | 0.6 |
| **Test Set III** | | | | | | | | | | | | | | | | | | |
| RASP | 1.09 | 0.36 | 1.29 | 0.45 | 1.33 | 0.42 | **2.36** | **0.64** | 1.05 | 0.32 | 1.45 | 0.37 | 1.02 | 0.45 | 1.13 | 0.37 | 1.63 | 0.41 |
| ϵSCORE | 2.54 | 0.37 | 3.14 | **0.53** | 2.71 | 0.42 | 1.31 | 0.46 | 2.47 | 0.4 | 2.74 | 0.41 | 2.59 | **0.53** | 2.75 | 0.46 | **3.44** | 0.47 |
| DFIRE-RNA | 3.37 | 0.51 | 3.47 | 0.53 | 3.68 | 0.54 | 2.06 | **0.58** | 3.2 | 0.45 | 3.58 | 0.46 | 3.96 | 0.55 | 3.5 | 0.43 | **4.39** | 0.51 |
| rsRNASP | 4.51 | 0.56 | 4.74 | 0.63 | 4.67 | 0.6 | 1.55 | 0.51 | 4.53 | 0.56 | 4.72 | 0.56 | 5.03 | **0.67** | 4.23 | 0.57 | **5.29** | 0.59 |

**Table 3.** ES and PCC scores for the three test sets for eight metrics. Different scales between datasets are explained by the different decoys' generation. Bold values mean the highest correlation for a given scoring function (for either ES or PCC).
